# Matched single-cell chromatin, transcriptome, and surface marker profiling captures in vivo epigenomic reprogramming during basal-to-luminal transition in the mammary gland

**DOI:** 10.64898/2026.02.16.706078

**Authors:** Anna Schwager, Eve Moutaux, Adeline Durand, Alexandra Van Keymeulen, Amélie Viaene, Mélanie Miranda, Louisa Hadj Abed, Simon Besson-Girard, Marion Lambault, Délia Dupré, Grégoire Jouault, Mélissa Saichi, Juliette Bertorello, Simon Dumas, Mathias Schwartz, Marthe Laisné, Justine Marsolier, Manuel Guthmann, Lorraine Bonneville, Urvashi Chitnavis, Déborah Bourc’his, Elisabetta Marangoni, Nicolas Servant, Cédric Blanpain, Leïla Perié, Céline Vallot

## Abstract

Single-cell multi-omics methods enable simultaneous mapping of chromatin states and transcriptomes, offering deep insights into gene regulation. Yet, the full potential of these approaches remains untapped for rare cell populations, as most methods require thousands of cells and are limited in their ability to capture multiple molecular layers comprehensively within the same cell. Here, we introduce OneCell CUT&Tag a user-friendly method that provides matched high-resolution epigenome, full-transcriptome, and surface marker quantification from every cell, with input as low as one cell. Using this approach, we uncovered epigenomic priming of basal cells in the mammary gland and captured the dynamics of basal-to-luminal transdifferentiation. We identified a transitional cell population with intermediate epigenomic profiles—absent in reference populations—and demonstrated a continuous epigenomic progression from basal to luminal states, while transcriptomes exhibited a binary switch. Adaptable to diverse samples and tissues, this method also revealed the role of H3K27me3 in shaping zygotic expression programs. By matching multiple layers of molecular information at single-cell resolution, OneCell CUT&Tag dissects the complementary roles of each omics layer in shaping cellular identity and function, opening new avenues to study rare and complex biological systems.

## Introduction

Histone modification profiling captures the diversity of chromatin states across both euchromatin and heterochromatin, providing insights beyond chromatin accessibility assays. For instance, these techniques have revealed how stem cells differentiate into specialized cell types^1–3^. In cancer research, single-cell chromatin profiling has revealed features of non-genetic tumor heterogeneity^4–6^ and mechanisms underlying drug resistance acquisition^7^. When integrated with RNA profiling, single-cell histone profiling sheds light on the dynamics of gene regulation and cell fate^8–10^, as demonstrated in mouse models.

Among the methods for single-cell histone profiling, a few are compatible with transcriptomic analysis of the same cells (Extended Data Fig. 1a). Yet, the vast majority require at least 10,000 cells as starting material, limiting their application to low-input samples such as *in vivo* or patient-derived material. This high input requirement stems from bulk antibody-based histone targeting prior to single-cell isolation (Extended Data Fig. 1b). Droplet-based and combinatorial indexing methods (e.g., CoTECH^11^, Paired-Tag^8^, scMTR-seq^12^), while suitable for high-throughput profiling, demand substantial input, profiling hundreds to thousands of cells from ≥100,000 starting cells. Lower-throughput, plate-based approaches (e.g., sortChIC^3^, T-ChIC^10^) reduce input requirements but still need ≥10,000 cells. Emerging low-input methods (e.g., itChIP^13^, TACIT-seq^14^) can profile <100 cells, but do not provide matched epigenomic and transcriptome data from the same cells (Extended Data Fig. 1a-b).

Here, we propose OneCell CUT&Tag which collects multiple layers of information on the same individual cell. OneCell CUT&Tag is a fast and user-friendly approach for chromatin profiling, transcriptomic and surface marker analysis starting from the same cell, adapted to multiple sample types. OneCell CUT&Tag enables the comparison of the different layers of omics information on individual cells.

## Results

### OneCell generates high-coverage epigenomes starting from individual cells

Compared to other existing low input epigenomic methods, the ambition of OneCell CUT&Tag was to perform the epigenomic profiling on individual nuclei from start to end, to enable the collection of matched epigenome, transcriptome and surface marker data on the same cells (Fig. 1a, Extended Data Fig. 1b-c). OneCell CUT&Tag method - here after named OneCell - starts from single isolated cells in plate making it compatible with very low input material (>=1 cell) and phenotype correlation using surface marker quantification. OneCell can profile from a single cell to up to ∼10^3^ cells using multiple 384-well plates in 1.5 days. Cells can be either sorted individually (e.g., via surface markers) or directly pipetted (e.g. from zygotes) into 384-well plates. For multi-omic characterization, a fraction of cytoplasmic mRNAs is pipetted to a mirror plate that will be processed with an adapted FLASH-seq protocol^15^ to produce full-length mRNAs sequencing libraries for each cell. Critically, the entire CUT&Tag part of the protocol —from nuclear isolation to sequencing library preparation— is performed on individual cells without the need for a transitory pool of cells (Fig. 1b, Extended Data Fig. 1b-c).

**Fig. 1:**
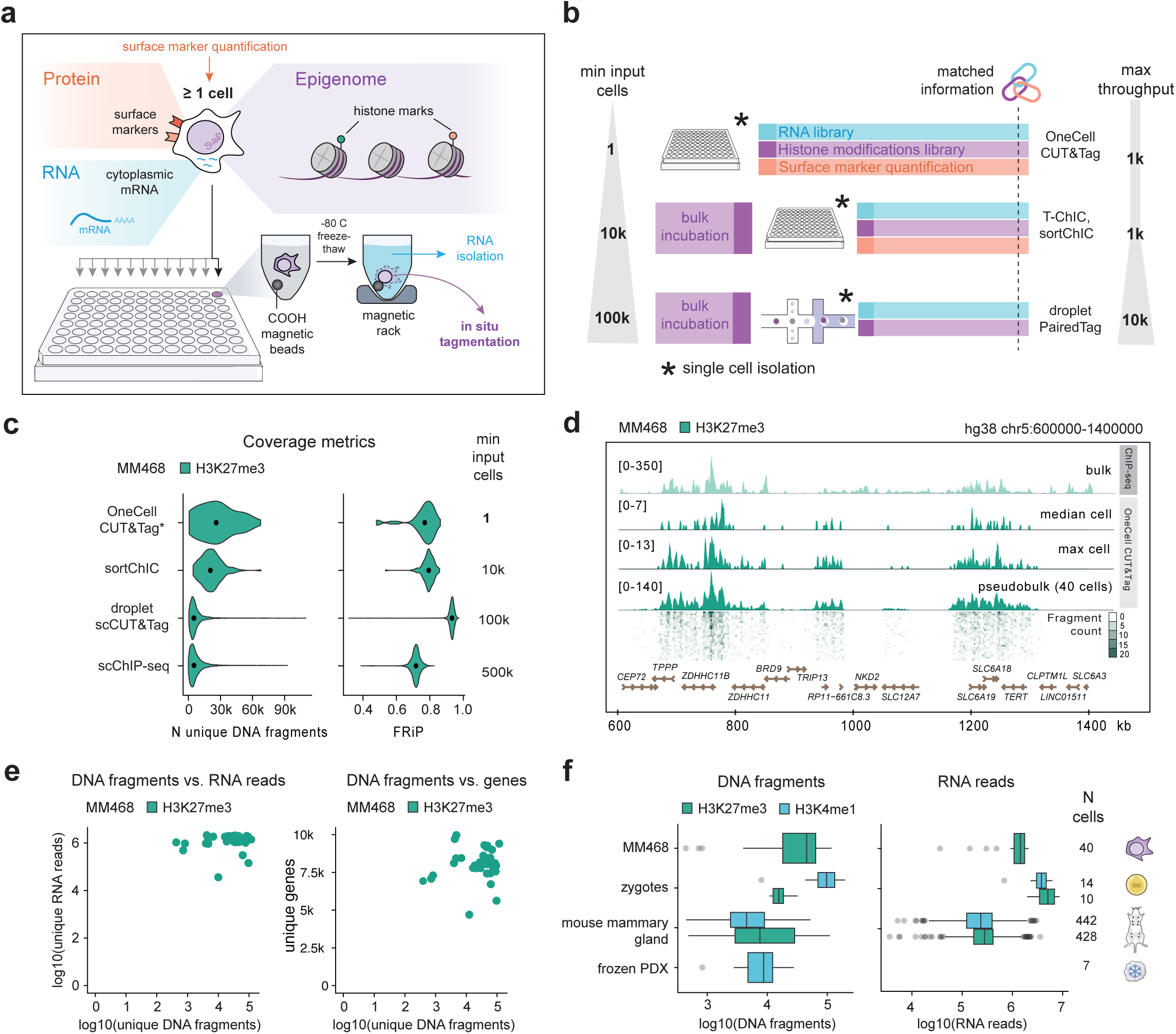
OneCell CUT&Tag workflow and benchmarking. **a.** Schematic representation of the OneCell CUT&Tag workflow and its integration with FACS phenotyping and RNA profiling. **b.** Comparative scheme of reference workflows allowing simultaneous transcriptomic and histone modifications profiling at the single cell level. **c.** Quality metrics for H3K27me3 profiling of MDA-MB-468 cells using four different single-cell chromatin profiling techniques: OneCell CUT&Tag, sortChIC, 10X droplet scCUT&Tag and scChIP-seq. Sequencing data from OneCell were bioinformatically subsampled to have similar sequencing coverage as other methods. The dot within violin plots indicates median values. **d.** Example of H3K27me3 coverage tracks obtained using OneCell CUT&Tag compared to bulk ChIP-seq in the MDA-MB-468 cell line. For OneCell CUT&Tag, pseudobulk (n=40 cells) and single-cell tracks from best-covered (118k fragments) and median-covered (45k fragments) cells are shown. **e.** Scatterplot representing pairwise metrics of unique DNA fragments (H3K27me3 profiling) and unique RNA reads or unique genes detected per cell for MDA-MB-468 cells (n=40 cells). **f.** Quality metrics for OneCell CUT&Tag performed on different sample types for two different histone marks: H3K27me3 (green) and H3K4me1 (blue) without subsampling. Boxplots show the median and interquartile range (Q1–Q3), whiskers correspond to 1.5x interquartile range, and outliers are shown as individual dots.

To obtain sufficient epigenomic data to interpret signals from individual cells, we optimized key protocol steps to minimize loss of nuclei and DNA fragments and enhance signal to noise ratio while limiting the number of processing steps (Extended Data Fig. 1c, Methods). We first fine tuned the composition of the lysis buffer, ensuring both chromatin integrity and capture of cytoplasmic mRNAs. Using carboxylic beads for nuclei isolation allowed serial add and removal of solutions without any loss of material. Adjustment of NP-40 detergent concentration in final nuclei lysis was instrumental to ensure fine nuclei lysis and chromatin accessibility without any DNA loss. Finally we adapted the FLASH-Seq protocol^15^ for transcriptome profiling - buffers and PCR cycles - to cytoplasmic extracts that were limited in material compared to whole-cell mRNAs extracts.

To evaluate the performances of OneCell for epigenomic profiling, we benchmarked it to reference methods using a breast cancer cell line MDA-MB-468 (Fig. 1c, Extended Data Figure 2, 3a). We focused on the repressive epigenomic mark H3K27me3, associated with facultative heterochromatin - a chromatin state that can only be analyzed by direct profiling of histone modifications. We selected two existing chromatin profiling methods that have multi-omic capacity – sortChIC^3^ and 10X droplet scCUT&Tag^1^. Both workflows are compatible with transcriptome profiling^8,12^, and rely on a respective minimum starting material of 10^4^ and 10^5^ cells. We also used as reference a scChIP-seq^5^ method that we had previously developed, chromatin immunoprecipitation being a method of choice for repressive chromatin profiling.

In order to compare approaches, we designed a scalable and reproducible computational workflow that can handle sequencing files from any single-cell epigenomic approach, provided it receives the whitelist of cell barcodes (Extended Data Fig. 2; see Methods). After equalizing the amount of sequencing reads per cell across all four methods (38k sequencing reads/cell in average) (Extended Data Fig. 3b-c), we observed that OneCell provided high-resolution epigenomes (Fig. 1c; 26,008 median unique reads/cell; 0.77 median fraction of read ends in peaks [FrIP],) starting from individual cells. Its performances were similar to the sortChiC method (20,581 median reads/cell; 0.79 median FrIP). Both methods provided a higher median coverage than high-throughput droplet methods, which in our hands had a median coverage of 5,722 (scChIP-seq) and 5,156 (10X droplet scCUT&Tag) unique reads per cell respectively. While high-throughput approaches produced sparse data that require pseudo-bulk visualization and analysis of clustered cells, with OneCell’s high-coverage approach, users can resolve single-cell coverage tracks at 1-Mb genomic resolution (Fig. 1d). Bulk ChIP-seq tracks analyzed from a million cells are shown as reference.

We then checked that high performances for epigenome profiling were compatible with high performances on matched transcriptomic profiling. We achieved for each cell extensive transcriptome coverage (median 8,054 genes/cell), while obtaining over 15k coverage for epigenome in 82% of cells (Fig. 1e, Extended Data Fig. 3d-e). These combined metrics on individual cells showed multi-omic capacity on each cell with no need for metacell pooling to achieve sufficient coverage for downstream analysis.

### OneCell provides matched multi-omic profiling for diverse sample types and supports automation

To assess its broader relevance and practicality —particularly for tissue samples beyond cell lines— we profiled repressive (H3K27me3) or permissive (H3K4me1) chromatin states in mouse and human tissues, both fresh and frozen (Fig. 1f). For fresh samples, we generated matched epigenomic and transcriptomic profiles, while for frozen samples, we isolated nuclei (not whole cells) and performed chromatin profiling alone. Performance metrics were highest for one-cell zygotes from embryos (15,590 median H3K27me3 DNA fragments/cell, 97,849 median H3K4me1 DNA fragments/cell, 13,712 median genes/cell) and the MDA-MB-468 cell line (46,486 median H3K27me3 DNA fragments/cell, 8,040 median genes/cell). As expected, human and mouse tissues showed lower metrics than the reference cell line MDA-MB-468 (e.g., 7,556 median H3K27me3 DNA fragments/cell and 2,404 genes/cell in the fresh mouse mammary gland). Notably, we also obtained individual epigenomes from a frozen human triple-negative breast cancer (TNBC) tumor (8,706 median H3K4me1 DNA fragments/cell), demonstrating OneCell’s versatility across sample types, in particular for patient derived material.

To enhance throughput and robustness, we automated the OneCell workflow using liquid-handling robots, replacing manual multi-channel pipetting. We progressively automated all dispensing and pipetting steps (Extended Data Fig. 4a), observing a 1.5X increase in single-cell coverage with full automation, with a maximum of 82k DNA fragments (H3K4me1) and 6k genes per cell (Extended Data Fig. 4b). Additionally, we found a correlation between the final concentration of amplified DNA/cDNA and the number of unique DNA fragments or genes per cell (Extended Data Fig. 4c-e), indicating that library quantification could serve as a cost-effective predictor of sequencing success. Overall, automation reduced hands-on-time enabling the processing of several plates in parallel, 4 vs 1 plate processed in 1.5 days (1,536 vs 384 cells).

### H3K27me3 shapes gene expression patterns in the zygote

To demonstrate OneCell’s capacity for multi-modal single-cell profiling, we applied our method to individual mouse zygotes (Fig. 2a). This enabled the first matched mapping of a zygote’s transcriptome alongside its repressive (H3K27me3) or permissive (H3K4me1) epigenome—both derived from the same cell. We generated 10 H3K27me3 and 14 H3K4me1 profiles from zygotes isolated from two mice, all paired with full-length transcriptomes. Epigenomic coverage exceeded 100 kb unique reads for some zygotes (Fig. 1f; Extended Data Fig. 5a-b). These profiles closely matched reference epigenomes, including bulk data we generated from pooled zygotes (n=50) and pseudo-bulk from published single-cell omic datasets^14^ (Fig. 2b-c; Extended Data Fig. 5c). High correlation between zygotes indicated minimal biological variation at this stage (Fig. 2d). While H3K4me1 accumulated genome-wide without clear enhancer or promoter association—typical of differentiated cells— H3K27me3 displayed a classical pattern, forming large domains spanning several megabases (Fig. 2b, e, f).

**Fig. 2:**
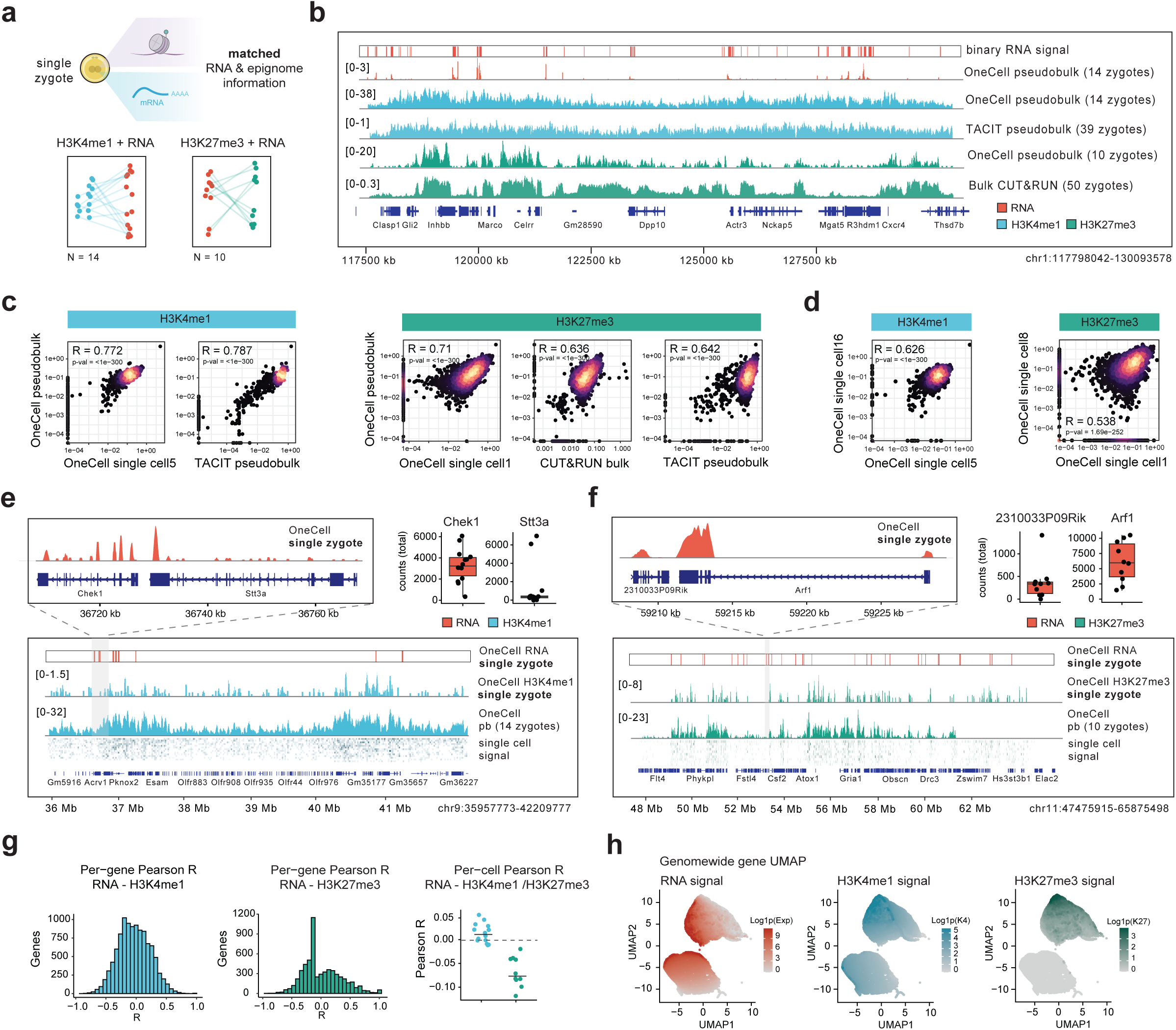
OneCell CUT&Tag provides matched transcriptome and epigenome profiling of single zygotes. **a.** Schematic representation of the multi-omic profiling performed for single mouse zygotes. Matched transcriptomic and epigenomic information was obtained for a total of 24 zygotes: 14 zygotes profiled for RNA and H3K4me1 mark, 10 zygotes profiled for RNA and H3K27me3 mark. Lines connect the same cell in each modality, matching epigenome profiles and transcriptomes for individual cells. **b.** H3K27me3 (green), H3K4me1 (blue) and RNA (red) profiles over a representative region of chromosome 1, obtained using OneCell CUT&Tag for 10 (H3K27me3) or 14 (H3K4me1) zygotes, or reference methods bulk CUT&Run and TACIT. For RNA profiles, a binary signal is shown alongside the coverage track to improve visibility, which is otherwise obscured by the large dynamic range of the data. **c.** Scatter plots showing correlations of H3K4me1 (blue) or H3K27me3 (green) signals between OneCell CUT&Tag pseudobulk profiles, OneCell CUT&Tag single-cell profiles, and reference methods (pseudobulk TACIT for H3K4me1 and bulk CUT&Run for H3K27me3). Dots correspond to 0.5 Mb genomic bins. Correlation scores and p-value from two-sided Spearman correlation test are indicated. **d.** Scatter plots showing correlations of H3K4me1 (blue) or H3K27me3 (green) signals over 0.5 Mb genomic bins between OneCell CUT&Tag single-cell profiles for two random cells. Correlation scores and p-value from Spearman R and the p-value of the two-sided Spearman correlation test are indicated. **e.** H3K4me1 (blue) and expression (red) profiles over a representative region on chromosome 9; pseudobulk for 14 zygotes and single zygote tracks are shown. Zoom-in displays whole-gene transcriptome coverage Boxplots show the expression level of corresponding genes (the box shows the median and interquartile range (Q1–Q3), the whiskers are 1.5x interquartile range, all individual points are shown). **f.** Same as (e) for H3K27me3 mark profiled in 10 single zygotes. **g.** Correlations between RNA expression (red) and H3K4me1 (blue)/H3K27me3 (green) signals measured in the same single cells. Chromatin signals were computed over the gene body + 5kb region upstream of the TSS. Left and middle panels show the distribution of per-gene Pearson correlation coefficients, computed across single cells within individual genes. The right panel shows per-cell Pearson correlation coefficients, computed across genes within individual cells. Each dot represents one single cell; horizontal bars indicate the mean. **h**. UMAP embedding based on genome-wide RNA expression, H3K4me1, and H3K27me3 signals. Each dot represents a gene. Color intensity indicates the signal strength for RNA expression (red), H3K4me1 (blue), or H3K27me3 (green).

We then explored the relationship between transcriptional activity and epigenomic landscapes. Leveraging matched RNA and epigenomic information for all zygotes, we compared gene expression levels and chromatin states across zygotes and within individual cells. Gene expression showed no correlation with H3K4me1 levels (Fig. 2g, left panel), consistent with the mark’s genome-wide, non-specific distribution in these cells. In contrast, we identified a subset of genes repressed by H3K27me3 (Fig. 2g, middle panel). At the single-cell level, H3K4me1 distribution remained unrelated to expression, while H3K27me3 emerged as a negative regulator of gene expression (Fig. 2g, right panel). Finally, a gene-based embedding analysis, integrating multi-modal data, revealed two distinct gene groups: both enriched for H3K4me1 but differing in H3K27me3 enrichment (Fig. 2h).

Taken together, these results showed that at this developmental stage, all genes appeared poised for activation, with H3K27me3 acting as a critical repressor—modulating transcription by targeting a specific gene subset.

### OneCell uncovers epigenomic priming of basal cells in the mammary gland

We used OneCell to study cell fate encoding in the mouse mammary gland, focusing on its epithelial compartment, which comprises basal and luminal lineages^16^. The luminal lineage further includes luminal hormone-sensing (ER+ LCs) and secretory precursor cells (ER- LCs)^16–18^. The basal lineage encompasses both basal progenitors and differentiated myoepithelial cells^16,19^ (Fig. 3a). Using cytometry, we distinguished basal and luminal lineages based on established surface markers (Fig. 3b; Extended Data Fig. 6a-b). We generated UMAP embeddings for RNA, H3K4me1, and H3K27me3 data (Fig. 3c; Extended Data Fig. 6c). Independent clustering of each omics layer (RNA, H3K4me1, H3K27me3) revealed basal-like and luminal-like identities, defined by gene expression or histone mark enrichment of marker genes (Fig. 3c-d, Extended Data Fig. 6d). Similarly to previous single-cell studies^16,20^, we identified three groups of mammary epithelial cells: a cluster of basal cells (basal), a cluster of luminal ER- cells (ER- LCs) and a cluster of luminal ER+ cells (ER+ LCs)^21^. Exploration of known marker genes demonstrated that marker gene expression aligns closely with epigenomic profiles (Fig. 3d; Extended Data Fig. 6e): Krt14 is for example expressed solely in basal cells with a specific H3K4me1 enrichment in this population, while it is repressed by H3K27me3 in the luminal lineage. We next explored whether we could refine the functional annotation of epithelial cells leveraging additional surface markers. To do so, we probed the cell surface marker Tspan8 previously identified as a basal stem cell marker^19^, enriched within a fraction of the basal cell population (Fig. 3a,e) Our analysis showed that Tspan8 is epigenomically regulated (Fig 3f-g): Tspan8-positive basal cells exhibited H3K4me1 enrichment at the gene promoter, while Tspan8-negative cells displayed higher H3K27me3 coverage, potentially repressing its expression (Fig 3g).

**Fig. 3:**
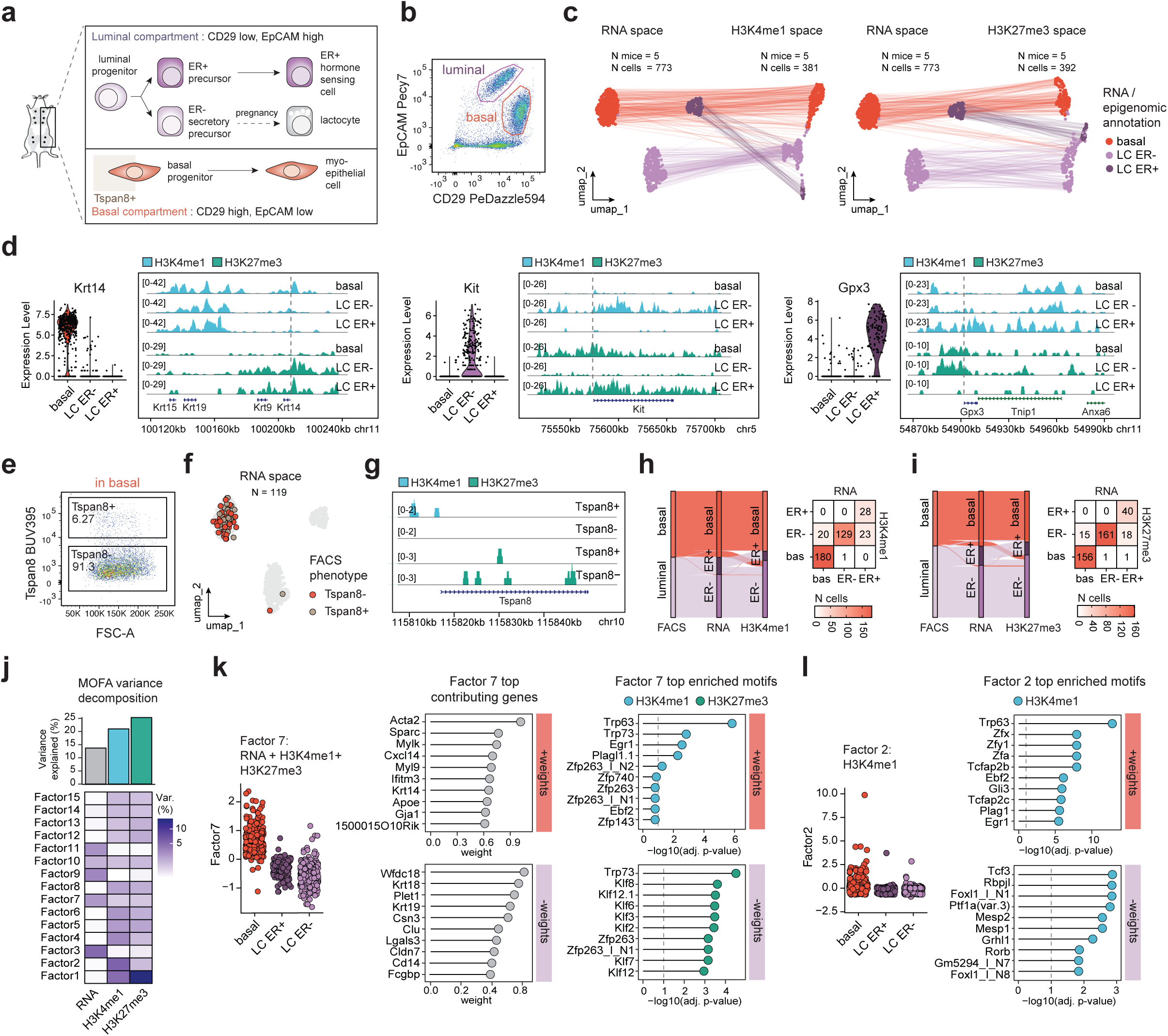
OneCell CUT&Tag multi-omic profiling decodes cell states in the mammary gland. **a.** Schematic representation of the two major epithelial lineages and cell types present in the adult mammary gland, together with the common cytometry surface markers used to distinguish between the lineages. **b.** Representative flow cytometry plot of EpCAM and CD29 surface marker expression used to identify basal and luminal epithelial cells. **c.** UMAP embeddings of individual cells generated separately for RNA, H3K4me1, and H3K27me3 modalities. Cells are colored according to unsupervised clusters identified within each modality and annotated based on cell-type marker gene expression or H3K4me1/H3K27me3 signal over marker genes. Lines connect representations of the same single cells across the different modalities. **d.** Expression of representative cell-type marker genes together with H3K4me1 and H3K27me3 coverage tracks across the gene bodies and surrounding genomic regions in the three mouse mammary gland cell populations. Dotted lines indicate the transcription start site (TSS) of the gene whose expression is shown in the violin plot. Pseudobulk H3K4me1 and H3K27me3 profiles are shown for each population. Cell-type annotations are based on the RNA modality. **e.** Representative flow cytometry plot of Tspan8 surface marker expression in the basal compartment. **f.** UMAP embedding generated from the RNA modality, with cells cytometrically profiled for Tspan8 surface marker expression highlighted. Cells classified as Tspan8-positive are shown in brown, cells classified as Tspan8-negative are shown in red. **g.** H3K4me1 and H3K27me3 profiles over the Tspan8 gene for the cytometry-defined Tspan8-positive and Tspan8-negative cells. **h.** Alluvial plots depicting concordance between cell-type annotations inferred from cytometry, RNA, and H3K4me1 data. To the right, the confusion matrix between the RNA- and H3K4me1-based annotations is shown. **i.** Same as (h) for the cytometry, RNA, and H3K27me3 concordance. **j.** Variance decomposition from a MOFA model jointly trained on RNA, H3K4me1, and H3K27me3 assays. Shades of violet indicate the proportion of variance explained by individual latent factors within each assay. Barplots show the cumulative proportion of total variance explained per assay. **k.** *Left:* Per-cell values of latent factor 7 from the joint MOFA model. Cell-type annotations are based on RNA modality. *Middle:* top genes contributing to Factor 7 from the RNA modality. The x-axis shows absolute feature loadings (weights). Features with positive weights are marked by a red bar, features with negative weights are marked by a violet bar. *Right:* top regulatory motifs enriched in genomic regions (5k bins) contributing to Factor 7 from H3K4me1 (blue) and H3K27me3 (green) modalities. The x-axis shows adjusted p-values of the motif enrichment analysis. Motifs enriched in features with positive weights are marked by a red bar, motifs enriched in features with negative weights are marked by a violet bar. **l.** Left: Per-cell values of latent factor 2. Cell-type annotations are based on the RNA modality. *Right:* top regulatory motifs enriched in genomic regions (5k bins) contributing to Factor 7 from the H3K4me1 modality. The x-axis shows adjusted p-values of the motif enrichment analysis. Motifs enriched in features with positive weights are marked by a red bar, motifs enriched in features with negative weights are marked by a violet bar.

We next assessed concordance between the three information layers by comparing cluster assignments of individual cells. Concordance between cytometry and RNA-based annotations for luminal and basal lineages was near-perfect (98% cells; Extended Data Fig. 6f). However, we identified discordances between RNA and epigenomic classifications of lineages for a fraction of cells (Fig. 3h-i). A subset of cells classified as basal by cytometry and RNA profiling exhibited luminal-like epigenomes—specifically ER- LCs (9% in average), in both H3K4me1 and H3K27me3 modalities. On the contrary, luminal cells almost never displayed basal-like epigenomes. Within the luminal lineage, classification of ER+ LCs and ER-LCs proved difficult, in particular for H3K27me3 because of continuity of the epigenomic data. ER- and ER+ LCs could be retaining epigenomic features of a common luminal progenitor (Fig. 3h-i).

To explore the unique contributions of each data layer, we integrated transcriptome and epigenome information using Multi-Omics Factor Analysis (MOFA)^22^ (Fig. 3j). This analysis identified factors specific to cell fates (Fig. 3k-l; Extended Data Fig. 6g-i). For example, factors 7 and 2 were basal-cell-specific, while factor 10 was ER+-specific (Extended Data Fig. 6h-i). Each factor’s contributing variance revealed the nature of its encoding: factor 2 was epigenome-specific, while factor 7 integrated multi-omic information (Fig 3j). Basal cell fate, encoded by factor 7, was characterized by actively transcribed genes (e.g., *Acta2*, *Krt14*) and basal transcription factor binding sites enriched for H3K4me1 (e.g., *Trp63*, *Trp73*; Fig. 3k). In contrast, H3K27me3 contributed to the repression of basal-associated transcription factor motifs in luminal cells (*Trp73* and *Klf* family binding sites). Factor 2 revealed an alternative H3K4me1 profile in a subset of basal cells, undetectable at the RNA level (Fig. 3l). Epigenomic enrichment was detected at genomic regions harboring motifs for stemness transcription factors, such as Zfx^20^, as well as transcription factors that drive both basal (Trp63) and luminal (Tcfap2c^23,24^) identity genes. These results highlight the epigenomic distinction of a basal cell subset, with epigenomic priming of stemness capacity, and underscore the value of integrating multi-omic data to decode complex cell fate encoding.

Overall, OneCell captured the added value of each omics layer to define cell fate. Our ability to uncover such benefits between omic layers stems from the method’s capacity to enable direct, cell-level comparisons, a resolution often unattainable with approaches relying on bioinformatic aggregation into metacells. Within the basal compartment, we detected epigenomic priming for luminal-like and stem-like features, undetectable at the RNA or protein level. This priming, specific to basal cells, aligns with their known multipotency in the adult upon lineage ablation^25^, transplantation^26–28^ or oncogenic Pik3ca^H1047R^ mutant expression^29^. In absence of such signals, we propose that this potential is epigenomically encoded in a fraction of basal cells - primed by H3K4me1, facilitating rapid luminal fate activation upon stimuli exposure.

### OneCell captures the epigenomic transition of basal to luminal lineage *in vivo*

To investigate the epigenomic dynamics underlying the basal-to-luminal fate transition, we used OneCell to profile rare cells during *in vivo* lineage conversion. To activate the multipotent differentiation potential of basal cells, we transplanted basal cells in the absence of luminal cells to mimic regenerative conditions^28^. We transplanted 10,000 fluorescent basal cells (isolated from K14cre/RosaYFP donor mice) into the cleared fat pads of immunodeficient Nod-Scid mice (Fig. 4a). We collected the rare engrafted cells at 4.5 days post-transplantation (n∼194 cells) and generated joint H3K4me1 and transcriptomic profiles for 46 of these cells. We also included profiling of the entire epithelial population from the K14cre/RosaYFP mouse (Fig. 4b; Extended Data Fig. 7a). By day 4.5, engrafted basal cells had reconstituted the epigenomic diversity of the epithelial compartment: 52% remained basal, 26% had transdifferentiated into ER- LCs, and 13% into ER+ LCs (Fig. 4b-c). Notably, leveraging the H3K4me1 diffusion map representation, we identified cells with intermediate epigenomic profiles—that are not found in the reference epithelial population—positioned between basal and ER- LCs branches (Fig. 4b).

**Fig. 4:**
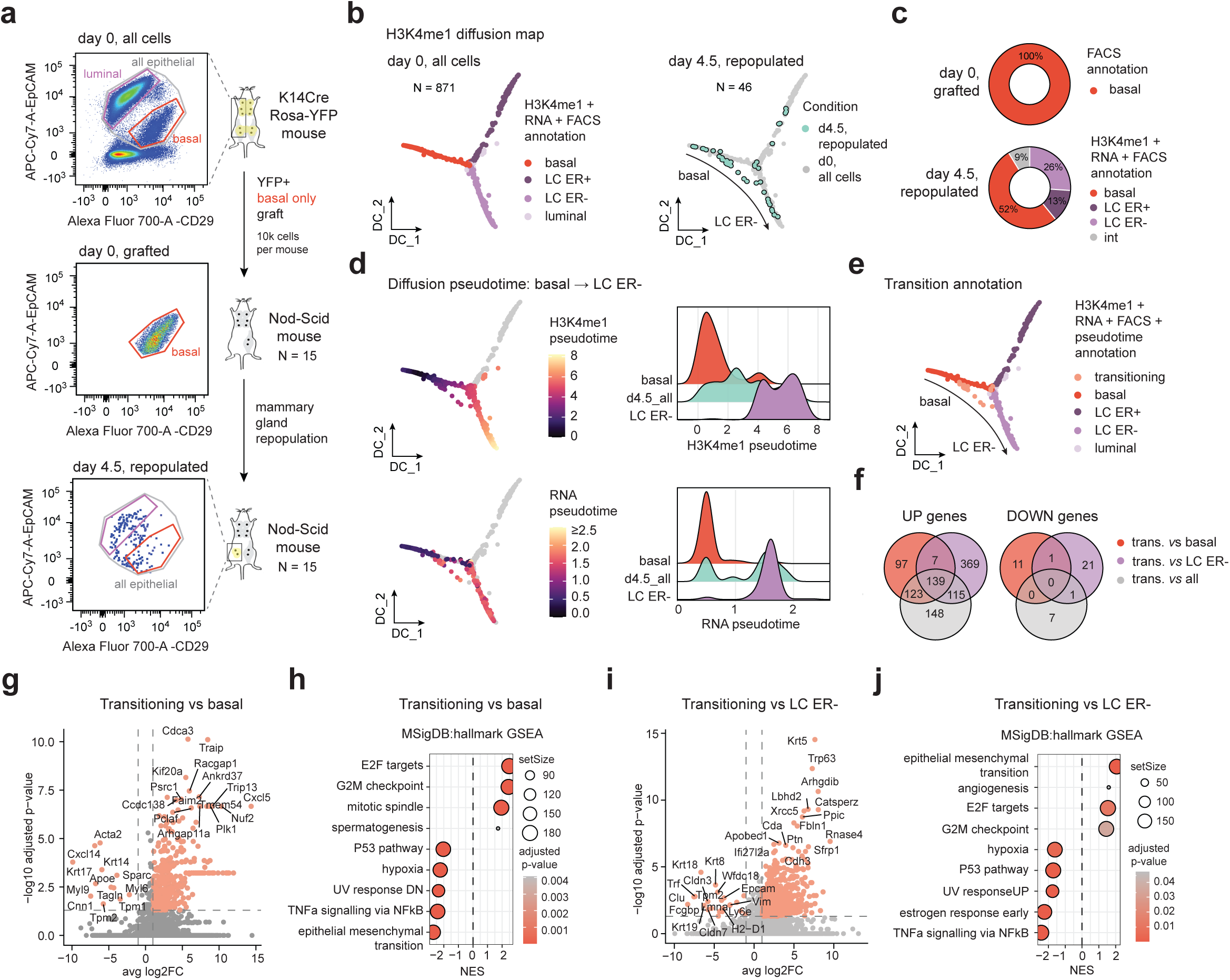
OneCell CUT&Tag captures the basal-to-luminal transition *in vivo.* **a.** Schematic representation of the experimental set-up and the representative flow cytometry plots of EpCAM and CD29 surface markers, defining basal and luminal lineages. **b.** Diffusion map constructed using H3K4me1 profiles from all cells highlighting: (*left)* reference control cells profiled at day 0 colored by cell-type annotations derived from combined H3K4me1, RNA, and cytometry data, and *(right)* epithelial YFP+ cells profiled at day 4.5. **c.** Cell type composition of engrafted cell population at d0 (based on cytometry data) and collected cell population at d4.5 (based on the joint H3K4me1, RNA and cytometry annotation). **d.** Diffusion pseudotime analysis capturing a basal-to–luminal trajectory. *Left:* H3K4me1 diffusion map colored by pseudotime values inferred from H3K4me1 signal (top) or inferred from RNA signal (bottom). *Right*: Density distributions of H3K4me1 (top) or RNA (bottom) pseudotime values for reference basal cells, reference ER- luminal cells, and cells collected at day 4.5. **e.** Representation of the transitioning cells defined by intermediate H3K4me1 pseudotime values on the reference H3K4me1 diffusion map. **f.** Venn diagrams showing the number of differentially upregulated (p-value_adj_ < 0.05, log₂FC > 1) or downregulated genes (p-value_adj_ < 0.05, log₂FC < −1) in transitioning cells compared with all other populations (grey), the basal population (red), or the ER- luminal population (violet). **g.** Volcano plot showing differentially expressed genes between transitioning and basal cells. Significantly deregulated genes (p-value_adj_ < 0.05, |log₂FC| > 1) are highlighted. The top 15 protein-coding genes (ranked by p-value_adj_) with |log₂FC| > 2 are indicated. **h.** Gene set enrichment analysis (GSEA) of genes differentially expressed between transitioning and basal cells using the mouse Hallmark gene sets from the MSigDB^32^. Top five positively and negatively enriched gene sets are shown. **i.** Same as (g) for the differentially expressed genes between transitioning and ER- luminal cells. **j.** Same as (h) for the genes differentially expressed genes between transitioning and luminal procursor cells.

To characterize these cells, we computed pseudotime trajectories based on H3K4me1 or RNA profiles. At day 4.5, descendants of engrafted basal cells exhibited a continuous epigenomic progression from basal to luminal lineage, while their transcriptomes showed a binary switch between basal and luminal identities (Fig. 4d). These results suggest a progressive epigenomic remodeling of basal cell transdifferentiating to luminal lineage, and rather a sharp transition from basal to luminal transcriptomes at the genome-wide level. We defined transitioning cells as the ones displaying intermediate pseudotime values, in between that of basal and ER- LCs (Fig. 4e). Comparison of the expression profiles of transitioning cells to basal or ER- LCs (Fig. 4f-j, Supplementary Table 1) revealed that cells undergoing epigenomic rewiring were more proliferative and exhibited downregulation of TNFα and p53 signaling. TNFα was actually shown previously to be one of the factors that restricts multipotency of basal cells *in vivo*^25^, validating our epigenomic identification of transdifferentiating cells. The temporary suppression of p53 signaling in these transitioning cells may facilitate cell fate conversion, as p53 is known to act as a barrier to cellular plasticity^30^. When compared to ER- LCs, transitioning cells also exhibited upregulated expression of receptor tyrosine kinase *Axl* reported to be a driver of stemness in the normal mammary gland^31^. Here, by integrating high-resolution multi-omic data, we further establish a connection between cell signaling mechanisms and the epigenomic regulatory programs driving genome rewiring during lineage conversion.

## Discussion

Using OneCell, we demonstrated how multi-modal profiling at high resolution uncovers previously unappreciated layers of regulatory complexity. In the basal compartment of the mammary gland, we compared, for each cell, different layers of complementary information, from surface markers, gene expression patterns to the regulatory epigenomic landscape. Histone modification profiling complemented transcriptome profiling, by providing additional refinements of the cell state encoding. A minority of basal cells displayed luminal-like epigenomes at the H3K4me1 and H3K27me3 levels. This priming was not detectable at the transcriptional or surface marker level, suggesting that epigenomic landscapes may precede and poise cells for future cell fate transitions, acting as an early regulatory layer. This observation aligns with the known context-specific multipotency of basal cells, which can regenerate luminal lineages but only upon lineage ablation or transplantation^26–28^. In physiological conditions, luminal and basal lineages are segregated, and do not interconvert^17,18^. We propose that H3K4me1-mediated priming, or H3K27me3 de-repression, of luminal-associated genes could establish an epigenomic "ready state," enabling rapid activation of luminal programs in response to environmental or developmental cues. Such a mechanism may underlie the adaptive plasticity of basal cells, ensuring tissue homeostasis and repair. We further grasped the molecular underpinnings of the transition from basal to luminal cell fate, through transplantation of basal cells in the absence of luminal cells. With OneCell, we show that the switch from basal to luminal cell fate happens through continuous epigenomic remodeling, but rather abrupt genome-wide transcriptomic changes.

Beyond the mammary gland, our zygote dataset serves as a starting resource for interrogating gene regulation principles during the earliest stages of mammalian development. By preserving single-cell resolution and enabling direct comparisons across omics layers, OneCell will potentially also be able to capture cell-to-cell variations as differentiation unfolds in the embryo. This paves the way for multi-modal mapping of further developmental transitions, offering insights into how regulatory programs are dynamically established.

Altogether, OneCell enables the direct matching and comparison of multiple omics layers at single-cell resolution, eliminating the need to bioinformatically aggregate cells into metacells to achieve sufficient coverage. The capacity to dissect the distinct contributions of each omics layer at the level of individual cells is essential for unraveling the molecular mechanisms underlying cell fate decisions. By enabling the decryption of regulatory transcription layers in challenging samples, notably *in vivo*, OneCell will facilitate the resolution of rare cell populations—decoupling the regulatory elements of expression patterns— particularly for patient-related material where cellular input is exceptionally low.

---

**Extended Data Fig. 1:**
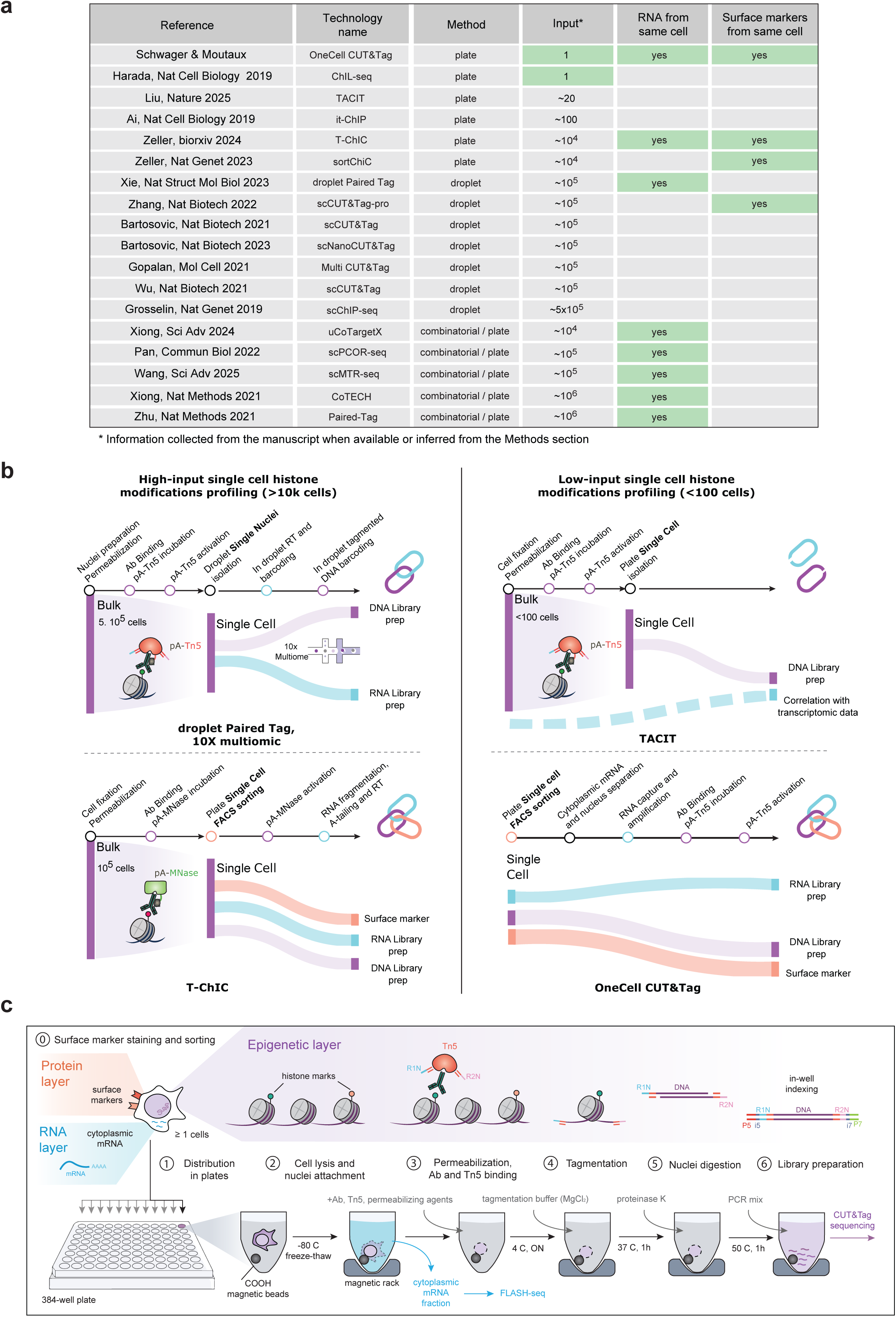
Detailed comparison of existing sc-Epigenomics methods. **a.** Comparative table of the existing methods capable of single cell histone modification profiling. **b.** Schematic representation of the OneCell CUT&Tag workflow and its integration with phenotype and RNA profiling. Detailed steps for the epigenomic part of the workflow are shown. **c.** Schematic overview of some single-cell epigenomic workflows, emphasizing the point of transition to single-cell isolation and how multi-omic measurements are integrated either experimentally or bioinformatically.

**Extended Data Fig. 2:**
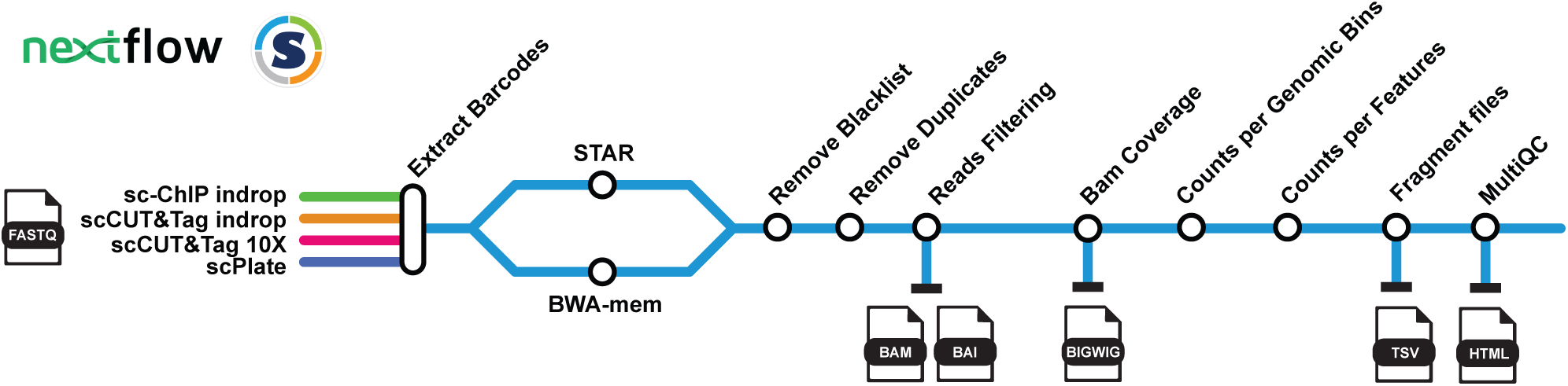
Reproducible and scalable bioinformatics pipeline for single-cell histone modification profiling. Input files are fastq sequencing files. The five output files of the workflow are depicted below the blue line.

**Extended Data Fig. 3:**
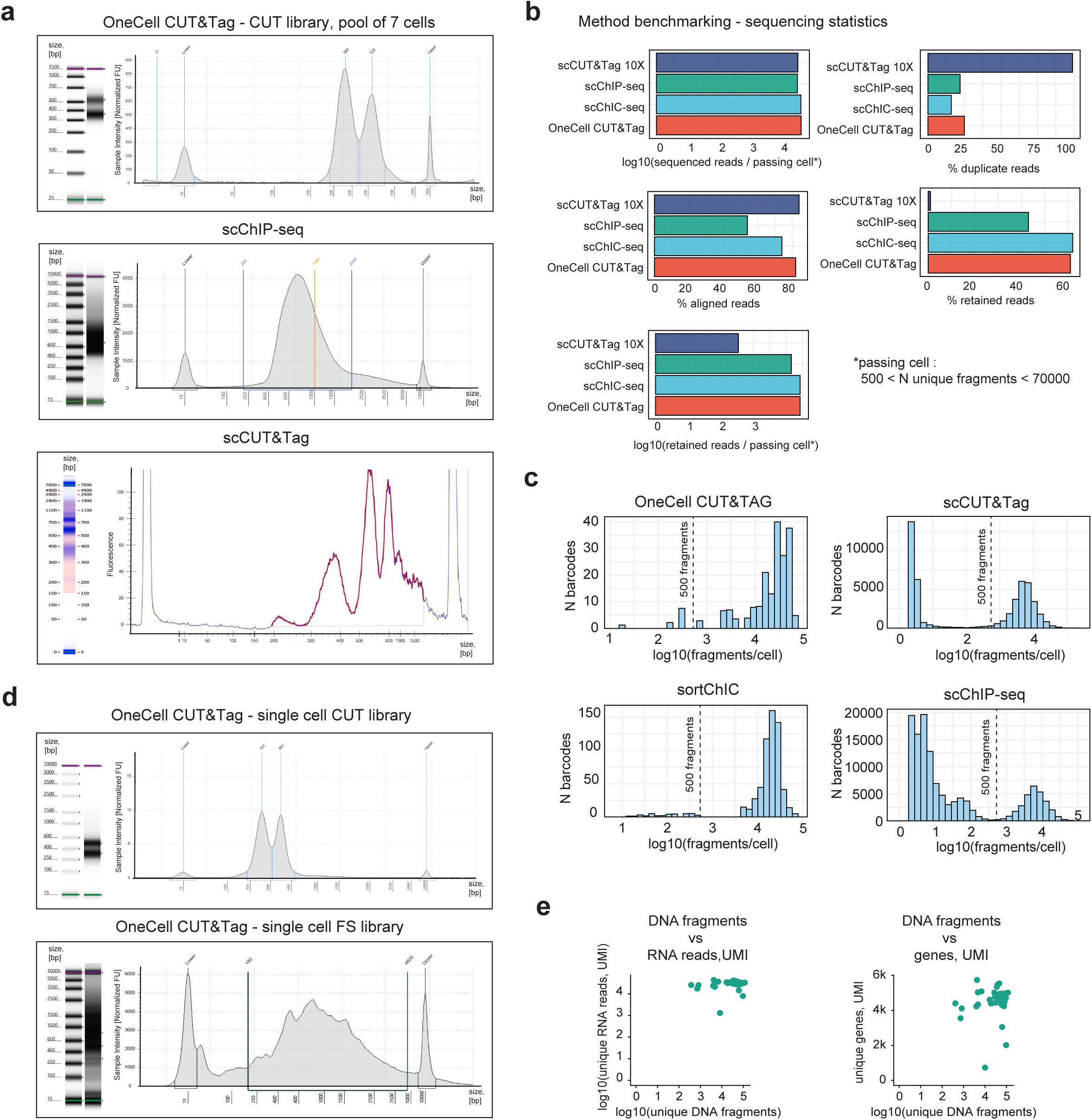
Metrics for the benchmarked methods. **a.** TapeStation profiles for benchmarked single-cell epigenomics techniques. **b.** Read statistics for benchmarked single-cell epigenomics techniques. **c.** Weighted histograms showing the numbers of unique DNA fragments detected per cell for four single-cell epigenomics methods: OneCell CUT&Tag, sortChiC, scChIP-seq and 10X droplet scCUT&Tag. The dashed lines show the filtering thresholds used to consider a cell as a “pass”. **d.** TapeStation profiles of CUT&Tag and FLASH libraries from a single cell profiled with OneCell CUT&Tag. **e.** Pairwise numbers of unique DNA fragments and unique RNA reads / unique genes based on the UMI-containing RNA reads only. Single cell data for each of the 40 MDA-MB-468 cells profiled for the transcriptome and the H3K27me3 mark.

**Extended Data Fig. 4:**
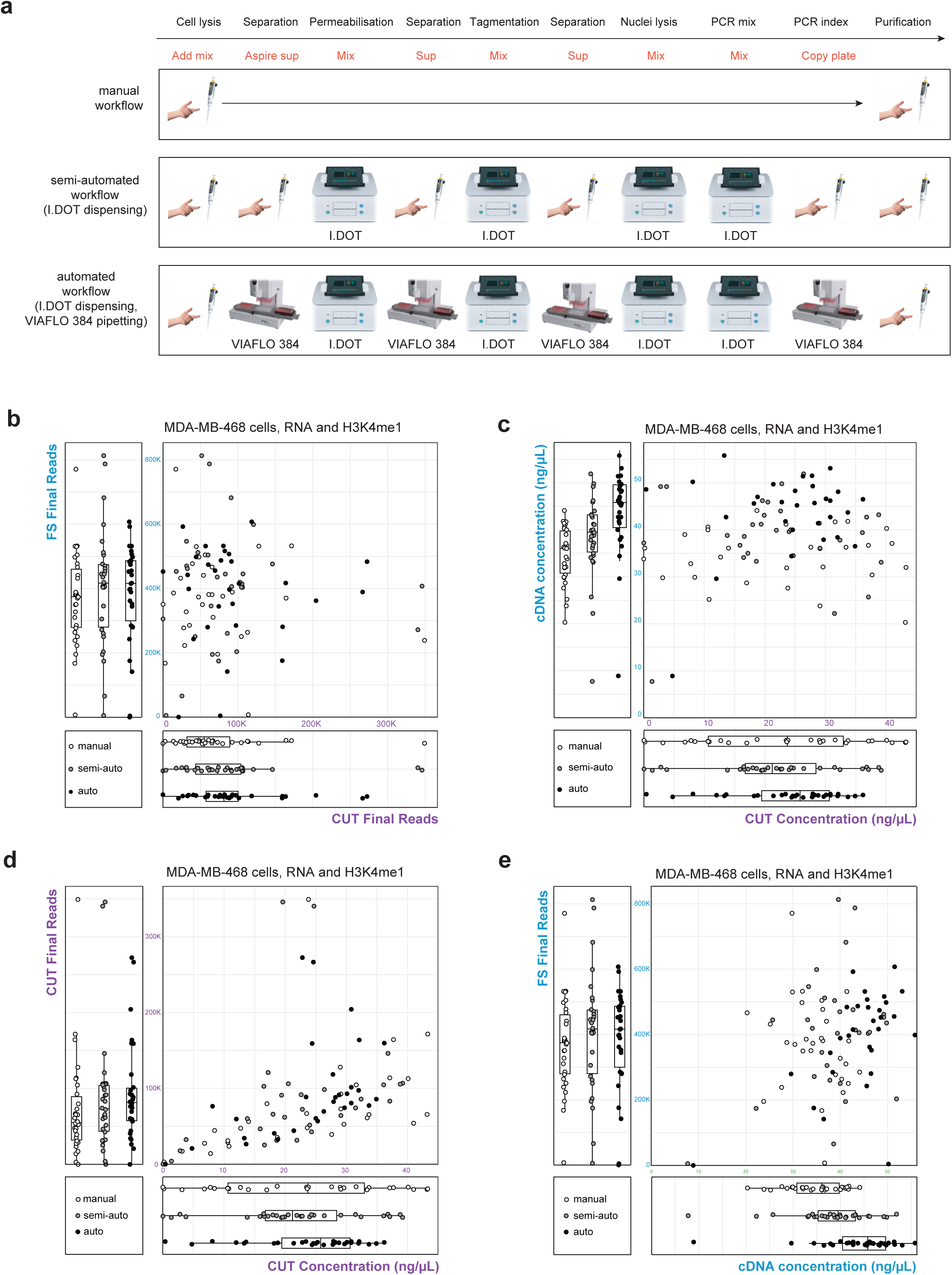
OneCell CUT&Tag automatisation. **a.** Schematic comparison of the manual, semi-automated and fully-automated OneCell CUT&Tag workflows. **b.** The number of final unique FLASH-seq reads (RNA modality, blue) and the number of final unique CUT&Tag fragments (DNA modality, violet) for the MDA-MB-468 cells profiled for the H3K4me1 mark and RNA using either the manual, the semi-automated or the automated OneCell CUT&Tag workflow. **c.** Scatterplot representing the concentrations of the cDNA versus DNA libraries for the MDA-MB-468 cells profiled for the H3K4me1 and RNA mark using either the manual, the semi-automated or the automated OneCell CUT&Tag workflow. **d.** Scatterplot representing the concentrations of the DNA libraries versus the final number of usable CUT&Tag reads for the MDA-MB-468 cells profiled for the H3K4me1 mark and RNA using either the manual, the semi-automated or the automated OneCell CUT&Tag workflow. **e.** Scatterplot representing the concentrations of the cDNA versus the final number of usable FLASH-seq reads for the MDA-MB-468 cells profiled for the H3K4me1 mark and RNA using either the manual, the semi-automated or the automated OneCell CUT&Tag workflow.

**Extended Data Fig. 5:**
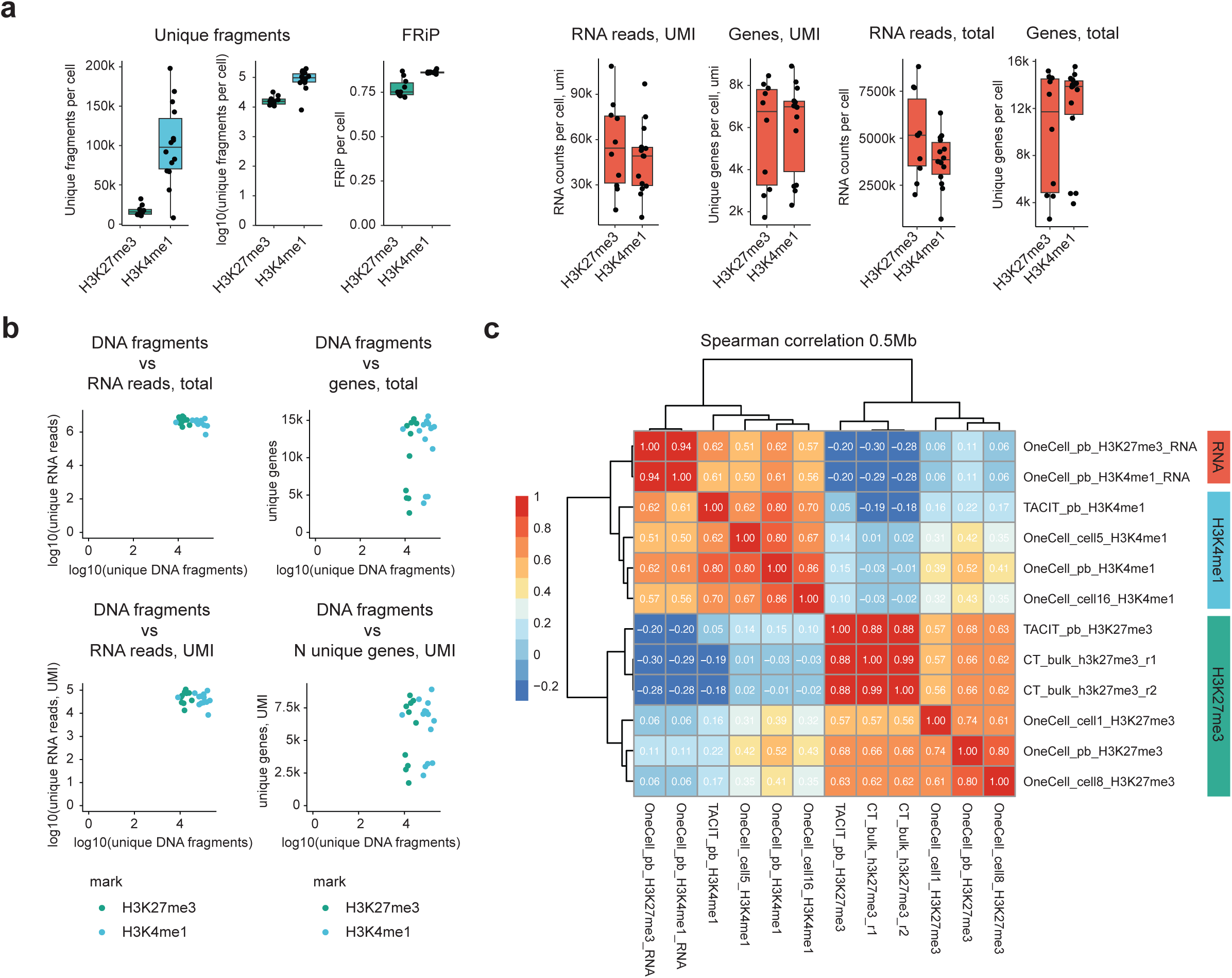
Multi-omic zygote profiling metrics. **a.** Quality control metrics for the multi-omic OneCell CUT&Tag experiment profiling RNA (red) with the H3K4me1 (blue) or the H3K27me3 (green) mark in single mouse zygotes. Boxplots show the median and interquartile range (Q1–Q3), the whiskers are 1.5x interquartile range; all individual points are shown. **b.** Pairwise numbers of unique DNA fragments and unique RNA reads / unique genes (total or UMI-only) for each of the 24 single zygotes profiled. The cells profiled for the RNA and H3K4me1 are colored blue, for the RNA and H3K27me - green. **c.** Spearman correlations between the RNA (red), H3K4me1 (blue) and H3K27me3 (green) signals over 0.5 Mb genomic bins obtained using OneCell CUT&Tag (pseudobulk and single cell profiles), TACIT (pseudobulk profiles) and bulk CUT&Tag.

**Extended Data Fig. 6:**
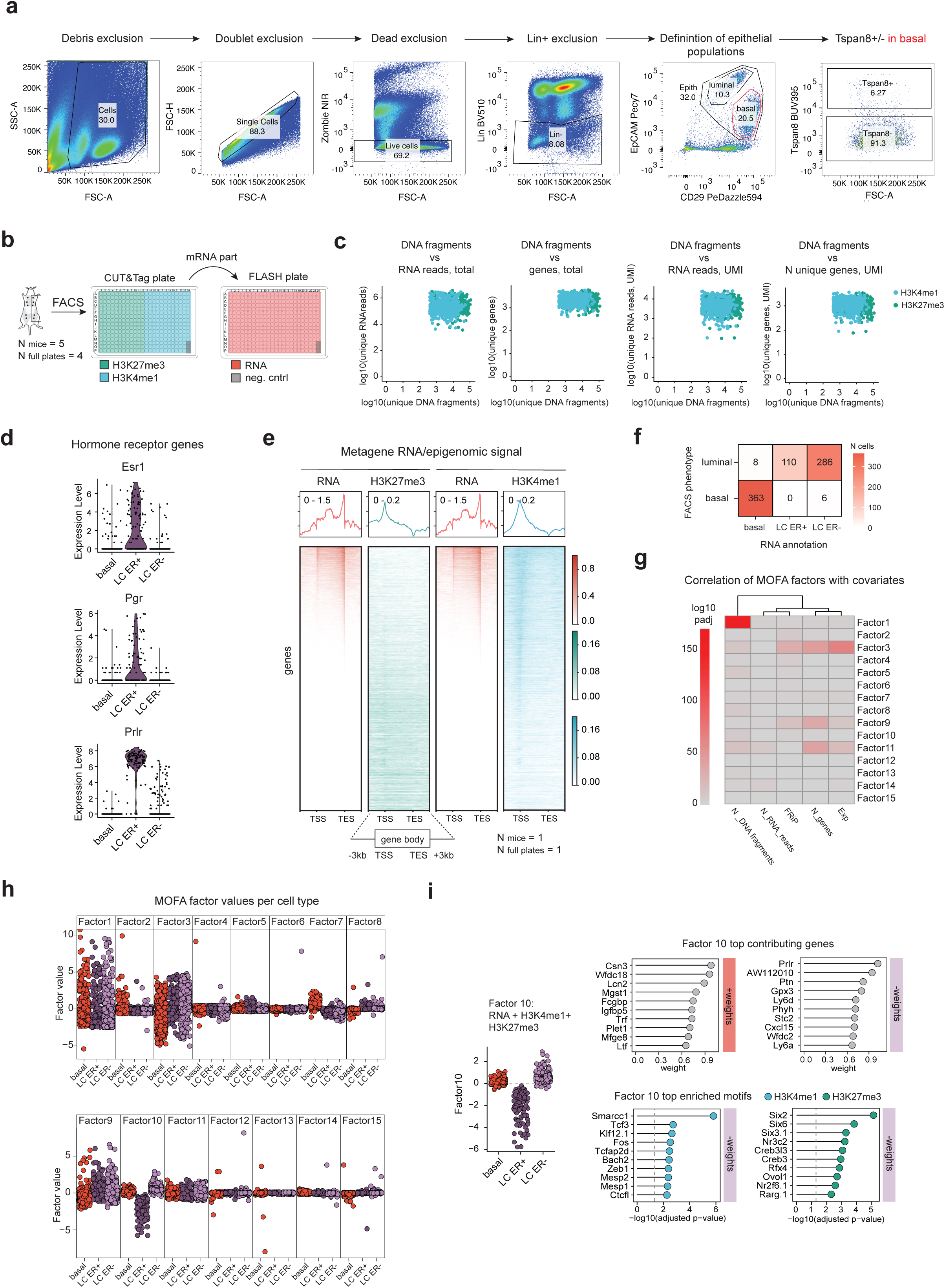
OneCell profiling of the mammary gland. **a.** Flow cytometry gating strategy used to sort the mouse mammary gland cells into OneCell CUT&Tag plates. **b.** Schematic representation of the plate design for the OneCell multi-omic profiling. **c.** Pairwise numbers of unique DNA fragments and unique RNA reads / unique genes (total or UMI-only) for each of the single cells profiled (N= 773). The cells profiled for the RNA and H3K4me1 are colored blue, for the RNA and H3K27me - green. **d.** Metagene representation of transcriptomic and epigenomic profiles across all coding genes. Data for one representative mouse replicate is shown. **e.** Confusion matrix showing the concordance between the cytometry-based and the RNA-based annotations. **f.** Correlation of quality control metrics and batch metadata with MOFA latent factors. Benjamini–Hochberg–corrected p-values from two-sided Pearson correlation tests are shown. Factors 1 and 3 are associated with technical variation. **g.** Per-cell values of all 15 latent factors captured by the joint RNA, H3K4me1 and H3K27me3 MOFA model. Cells are grouped by the annotation based on the RNA modality. **h.** *Left:* Per-cell values of latent factor 10 from the joint MOFA model. Cell-type annotations are based on RNA modality. *Middle:* top genes contributing to Factor 10 from the RNA modality. The x-axis shows absolute feature loadings (weights). Features with positive weights are marked by a red bar, features with negative weights are marked by a violet bar. *Right:* top regulatory motifs enriched in H3K4me1 (blue) or H3K27me3 (green) modalities. The x-axis shows adjusted p-values of the motif enrichment analysis. Motifs enriched in features with negative weights are shown.

**Extended Data Fig. 7:**
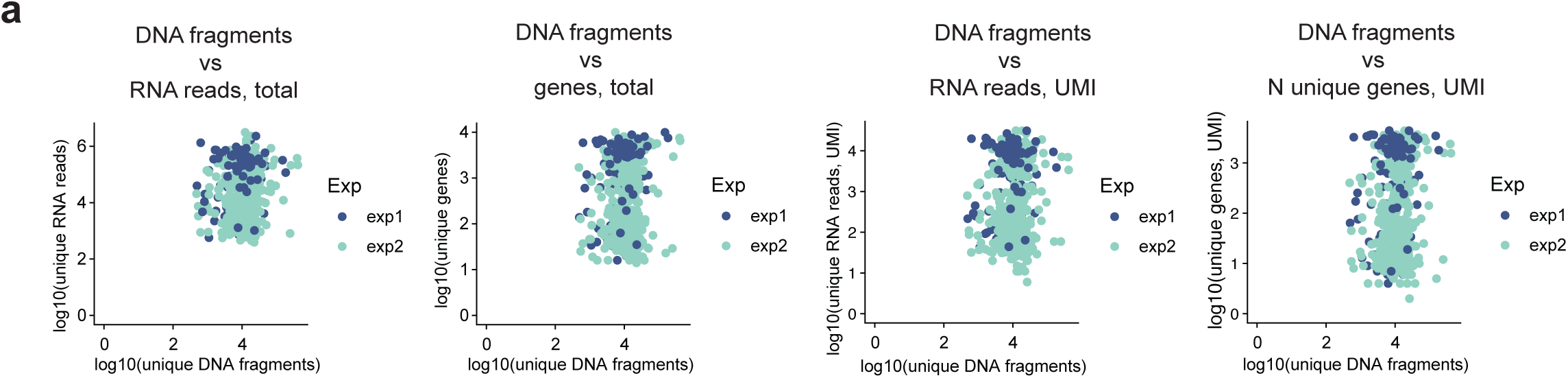
Related to the transplantation experiments. **a.** Pairwise numbers of unique DNA fragments and unique RNA reads / unique genes (total or UMI-only) for each of the single cells profiled (N = 545). All cells were profiled for the RNA and H3K4me1 and are colored according to the experiment replicate.

